# Intra-leaf modeling of *Cannabis* leaflet shape produces leaf models that predict genetic and developmental identities

**DOI:** 10.1101/2023.08.15.553356

**Authors:** Manica Balant, Teresa Garnatje, Daniel Vitales, Oriane Hidalgo, Daniel H. Chitwood

## Abstract

- The iconic, palmately compound leaves of *Cannabis* have attracted significant attention in the past. However, investigations into the genetic basis of leaf shape or its connections to phytochemical composition have yielded inconclusive results. This is partly due to prominent changes in leaflet number within a single plant during development, which has so far prevented the proper use of common morphometric techniques.
- Here we present a new method that overcomes the challenge of nonhomologous landmarks in palmate, pinnate and lobed leaves, using *Cannabis* as an example. We model corresponding pseudo-landmarks for each leaflet as angle-radius coordinates and model them as a function of leaflet to create continuous polynomial models, bypassing the problems associated with variable number of leaflets between leaves.
- We analyze 341 leaves from 24 individuals from nine *Cannabis* accessions. Using 3,591 pseudo-landmarks in modeled leaves, we accurately predict accession identity, leaflet number, and relative node number.
- Intra-leaf modeling offers a rapid, cost-effective means of identifying *Cannabis* accessions, making it a valuable tool for future taxonomic studies, cultivar recognition, and possibly chemical content analysis and sex identification, in addition to permitting the morphometric analysis of leaves in any species with variable numbers of leaflets or lobes.

## INTRODUCTION

*Cannabis sativa* L. (hereafter referred to as *Cannabis*) is a versatile crop plant used by humans for a variety of purposes throughout history. Although today it is commonly associated with its psychoactive properties, traditional medicine has relied heavily on *Cannabis,* and it is also a valuable source of food and fibers (Clarke & Merlin, 2013). Genetic and archaeological evidence suggests that *Cannabis* was domesticated around 12,000 years ago in East Asia, initially serving as a multipurpose crop before separate selections for fiber and drug production emerged around 4,000 years ago (Ren *et al*., 2021). Since then, widespread cultivation has facilitated its global distribution. Throughout the 20th century, *Cannabis* use was largely abandoned due to its illegal status in many parts of the world. However, recent legalization for recreational and/or medicinal purposes in many countries worldwide has led to a surge in the cannabis industry (*The Global Cannabis Report, 3rd Edition*, 2022).

Extensive *Cannabis* use has resulted in the development of numerous cultivars and strains that are well-suited to diverse uses and climates (Small, 2015). This significant morphological and phytochemical diversity within the *Cannabis* genus poses challenges for taxonomic classification. Over the past two centuries, various taxonomic approaches based on genetics, morphology, and phytochemistry have been proposed (McPartland & Small, 2020). Some scientists advocated for a polytypic classification, recognizing the presence of two (Lamarck & Poiret, 1783; Zhukovskii, 1971; Hillig, 2005a) or three (Emboden, 1974; Schultes *et al*., 1974; Hillig, 2005b; Clarke & Merlin, 2013) species with multiple subspecies, while others argued for a monotypic genus, considering only a single species, *Cannabis sativa* (Small & Cronquist, 1976; Sawler *et al*., 2015; Small, 2015; McPartland, 2018; McPartland & Small, 2020; Ren *et al*., 2021). Hillig (2005a) introduced a classification system based on biotypes, considering molecular, morphological, and phytochemical data. He proposed dividing *Cannabis* into two species, *C. sativa* and *C. indica* Lam., and six biotypes: *C. indica* as narrow-leaflet drug (NLD), wide-leaflet drug (WLD), hemp and feral biotype, and *C. sativa* as hemp and feral biotype. Recently, Lapierre *et al*. (2023) conducted a comprehensive taxonomic review of the *Cannabis* genus and based on available genetic data, strongly supported the theory that *Cannabis* is a highly diverse monotypic species.

Apart from taxonomic classification, *Cannabis* is often categorized based on its cultivation purpose, morphology, and chemical composition. Fiber-type plants, commonly known as hemp, are primarily grown for fiber and seed production. These plants contain less than 0.3% of the psychoactive compound THC (Δ9-tetrahydrocannabinol), while drug-type plants, often referred to as marijuana and medicinal cannabis, can contain higher levels of THC (Hurgobin *et al*., 2021). *Cannabis* plants can also be separated based on the ratio of two major cannabinoids THC and CBD (cannabidiol) into Type I (THC dominant), Type II (balanced CBD/THC ratio), and Type III plants (CBD dominant) (Small & Beckstead, 1973). In the medicinal and recreational cannabis industries plants are normally categorized as ‘sativa’, ‘indica’, or ‘hybrid’. Taller plants with narrow leaflets and high THC percentage are called ‘sativa’, while shorter and bushier plants with wider leaflets and high percentages of both CBD and THC are called ‘indica’. Plants with intermediate characters are called ‘hybrids’ (McPartland & Guy, 2017). While the classification of *Cannabis* into ‘indica’ and ‘sativa’ is not supported by genetic data, the visible differences in leaflet width have long been a significant characteristic used to visually discriminate different types of *Cannabis. Cannabis* arguably possesses one of the most iconic leaves among all plants. Its palmately compound leaves with a varying number of leaflets are a popular culture symbol. *Cannabis* exhibits a remarkable degree of phenotypic plasticity, further accentuated by selection pressure during the domestication process (Small, 2015). Extensive variability in leaf morphology has already been described by Quimby *et al*. (1973) and later Anderson (1980), who was the first to quantify the width, length, and ratio of the central leaflet. This or similar methods were then commonly used in studies investigating the morphological characteristics of *Cannabis* species, subspecies, cultivars, biotypes and chemotypes (Small *et al*., 1976; de Meijer *et al*., 1992; de Meijer & Keizer, 1996; Hillig, 2005a; Clarke & Merlin, 2013; Lynch *et al*., 2016; Karlov *et al*., 2017; Parsons *et al*., 2019; McPartland & Small, 2020; Carlson *et al*., 2021; Islam *et al*., 2021; Jin *et al*., 2021a; Vergara *et al*., 2021; Buzna & Sala, 2022; Chen *et al*., 2022; Murovec *et al*., 2022), often with contradictory results. Leaf shape has therefore played an important and sometimes controversial role in *Cannabis* taxonomy. While researchers in previous *Cannabis* studies were aware of enormous plasticity and the effect the environment has on leaf shape (Vergara *et al*., 2021; Murovec *et al*., 2022), they very rarely paid attention to the effects of developmental processes, even though heteroblastic changes (differences in leaf shape arising from juvenile-to-adult phase transitions in the meristem) profoundly affect the arrangement and shape of *Cannabis* leaves along the shoot. While some studies briefly mention the developmental changes of leaves (Hillig, 2005a; Carlson *et al*., 2021; Jin *et al*., 2021b; Spitzer-Rimon *et al*., 2022), the only two studies focusing on heteroblastic phase changes in leaves along the plant axis were done by Heslop-Harrison and Heslop-Harrison (1958) and Hesami *et al*. (2023). In the lower part of the shoot *Cannabis* leaves exhibit opposite phyllotaxy and one to three leaflets, transitioning to alternate phyllotaxy and leaves with up to 11 or 13 leaflets in the upper section (Hillig, 2005a; Clarke & Merlin, 2013; Small, 2015). Additionally, the changes in leaflet number are not uniform between different *Cannabis* accessions (Hillig, 2005a). These changes during development not only complicate categorization of plant accessions based on leaf shape, but also prevent the use of morphometric techniques.

Morphometrics is the quantitative analysis of shape. It includes a wide range of methods, from measuring allometric differences in dimensions like lengths, widths, and angles in relation to size (Niklas, 1994), to geometric techniques that measure shape comprehensively, like elliptical Fourier (EFDs; Kuhl & Giardina, 1982) and landmark-based analyses (Bookstein, 1997). It can be used to classify species and to separate effects on shape arising from genetic, developmental, and environmental mechanisms (Chitwood & Sinha, 2016). Historically the field of ampelography (ἄμπελος, ‘vine’ + γράφος, ‘writing’; Ravaz, 1902; Galet, 1952; Galet & trans. Morton, 1979) relied heavily on leaf shape to distinguish grapevine varieties. Unlike *Cannabis,* grapevine leaves have a consistent number of lobes, sinuses, and other associated homologous points that can be used for both landmark-based and EFD morphometric analysis (Chitwood *et al*., 2014; Chitwood, 2021) to disentangle genetic (Demmings *et al*., 2019), developmental (Chitwood *et al*., 2016a; Bryson *et al*., 2020; Migicovsky *et al*., 2022), and environmental effects (Chitwood *et al*., 2016b, 2021) embedded in leaf shapes.

The variable number of leaflets in *Cannabis* (and several other species with lobed, pinnate and palmate compound leaves) precludes analysis methods that rely on homologous, comparable points to measure shape comprehensively. Methods to automatically isolate individual leaflets (Failmezger *et al*., 2018) or to model developmental trajectories, such as heteroblastic series (Biot *et al*., 2016) were proposed previously for morphometrical analysis in such cases. In *Cannabis,* Vergara *et al*. (2021) used a landmark-based approach but were limited to analyzing the central and two most distal leaflets on each side, features that all *Cannabis* leaves except single-leaflet leaves possess, but which excludes most of the shape variation within a leaf.

Here, we seek to build on these works and conceptually extend our framework of continuously modeling leaflets within a palmate leaf. We model corresponding pseudo-landmarks for each leaflet as angle-radius coordinates relative to the petiolar junction and model angle and radius as a function of leaflet number to create continuous polynomial models that bypass the problems associated with variable numbers of leaflets between leaves. This enabled us to compare leaves with different numbers of leaflets within a plant and to discern differences between genotypes rather than the heteroblastic series. Analyzing over 300 *Cannabis* leaves, we model theoretical leaves with nine leaflets and 3,591 comparable pseudo-landmarks. Linear discriminant analysis (LDA) predicts accession, leaflet number, and relative node number with high accuracy. Intra-leaf modeling allows the application of morphometric techniques to comprehensively measure leaf shape in *Cannabis*, enabling future taxonomic and developmental studies, cultivar recognition, and possibly chemical content analysis and sex identification, in addition to permitting the morphometric analysis of leaves in any species with variable numbers of leaflets or lobes.

## MATERIAL AND METHODS

### Plant material and growing conditions

This study includes 24 individuals from nine accessions of *Cannabis sativa* L. (Table **1**; Fig. **1**), encompassing both wild/feral accessions and cultivated varieties with a wide distribution area. The plants were grown from seeds in a growth chamber (Fitoclima D1200PLL, Aralab, Portugal) to minimize the influence of the environment. Before sowing, the seeds were sterilized overnight in a 5% H_2_O_2_ solution with the addition of Inex-A solution (Cosmocel, Spain) at room temperature. Sterilized seeds were then transferred to Petri dishes and placed in the growth chamber for germination. Once the first leaves emerged, the seedlings were transferred to small peat pots with a pre-fertilized soil substrate (Kilomix Atami, Spain). During this phase, the environmental conditions were set to 25°C, with an 18-hour day and 6-hour night photoperiod, and a light intensity of 50 µmol m^−2^s^−1^ (Philips Master PL-L 55W, Spain). After two weeks the surviving plants were transplanted to 3.5 l pots with the same soil substrate. The light intensity was gradually increased to 300 µmol m^−2^s^−1^ over the following week, without changing the photoperiod and temperature. The onset of flowering in some *Cannabis* accessions is photoperiod dependent, therefore after four weeks, the photoperiod was changed to 12 hours of daylight and 12 hours of darkness, and the light intensity was gradually increased to 700 µmol m^−2^s^−1^ over the following week, while keeping the temperature at 25°C. The plants remained in these environmental conditions until the flowering stage. Plants received daily irrigation with tap water, without any application of nutrient or phytosanitary control.

**Table 1.** Accession details and number of leaves collected and analyzed in the study.

| Accession ID | Accession type | Location/Cultivar name | Number of individuals | Number of leaves collected | Number of leaves analyzed |
| --- | --- | --- | --- | --- | --- |
| AM15 | Wild/feral | Armenia, Sjunik marz, Goris town | 5 | 90 | 74 |
| BNG | Wild/feral | Bangladesh, Rangpur, Carmichael College Campus | 1 | 14 | 10 |
| FUT75 | Cultivar | Futura 75 | 2 | 45 | 30 |
| HU1 | Wild/feral | Hungary, Nyírvasvári | 4 | 83 | 68 |
| IK | Landrace | India, Kerala | 4 | 92 | 53 |
| IKL | Landrace | India, Kullu | 4 | 69 | 47 |
| MAR | Landrace | Morocco, North Morocco | 1 | 18 | 15 |
| MN9 | Wild/feral | Mongolia, Selenge aimag, Baruunburen sum | 1 | 14 | 10 |
| RO1 | Wild/feral | Romania, Mangalija | 2 | 36 | 34 |

**Fig. 1.**
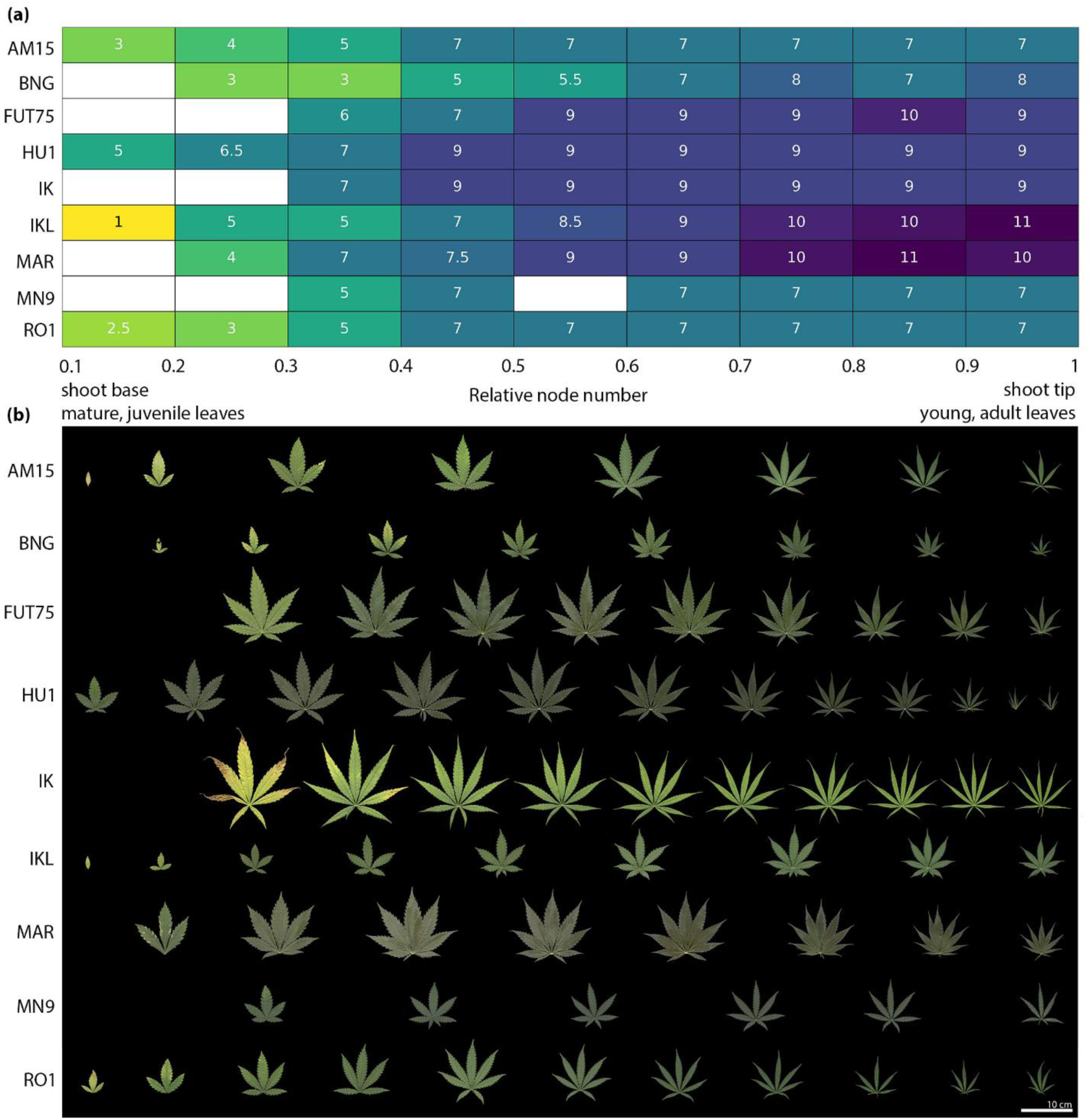
Changes in the leaf shape and leaflet number during the development in nine *Cannabis* accessions. (a) Median values for all available leaflet number for each relative node number for the nine *Cannabis* accessions. (b) Changes in leaf shape between different developmental stages in different *Cannabis* accessions.

### Leaf sampling and imaging

A total of 461 leaves were sampled during the flowering stage, with the exception of individuals from the accession IK, which did not begin to flower during the two-month cultivation period. Leaves along the main axis of the plants were collected and immediately scanned using a flatbed photograph scanner (Epson Perfection V370, Japan) at 1200 dpi resolution. A piece of velvet fabric was placed between the leaf and the scanner cover to avoid any shadows. No adjustments to the angle of individual leaflets were made before scanning. Each leaf was scanned with a scale and a label indicating the node it originated from, followed by a sequential lowercase letter, since typically two leaves are present per node. Starting at the base of the plant, the first two leaves were labeled as leaves “a” and “b” from node number 1, and so on, until the shoot apex.

*Cannabis* leaves display a marked heteroblastic, or juvenile-to-adult, leaf shape progression. Mature, juvenile leaves located on the first node at the base of the plant usually have a simple, serrated leaf. As node number increases so does the leaflet number, reaching a maximum of 9 to 13 leaflets in young, adult leaves at the growing tip. Eventually leaves transition into an inflorescence type. During this transition, the number of leaflets per leaf starts to decrease again until the top of the inflorescence. Leaves at the shoot base have opposite phyllotaxy and transition to alternate phyllotaxy in the upper section on the stem and inflorescence (Heslop-Harrison & Heslop-Harrison, 1958; Hillig, 2004; Potter, 2009; Spitzer-Rimon *et al*., 2022). To ensure that only stem leaves were included in our analysis, we separated the two types (i.e., stem and inflorescence leaves) based on the point where the decrease in the number of leaflets appeared. This point determined the “total node number”, the number of nodes per plant used for further analysis. Total node number varied among individuals. To compare node positions, a relative node number was calculated, which was defined by the node position divided by the total node number for the individual plant, where zero is at the plant base and one at the last node included in the analysis (Fig. **1**). Because of the nature of plant growth, the leaves at the base of the plant were frequently too senesced to be incorporated in the analysis or were entirely lost. Nevertheless, the nodes could still be identified, which allowed them to be taken into account in the calculation of relative node number.

### Image analysis and landmarking

After eliminating damaged and deformed leaves (39), simple leaves (4), leaves with even leaflet numbers (3) and leaves with relative node values above one (57), a total of 358 *Cannabis* leaves were used for image analysis and landmarking. Photoshop was used to separate petioles and leaflets smaller than 1 cm from the rest of the leaf. The leaf outlines were then extracted and saved using Python modules NumPy (Harris *et al*., 2020), Matplotlib (Hunter, 2007) and OpenCV (Bradski, 2000). The code for extracting and plotting the leaf outlines can be found on GitHub (https://github.com/BalantM/Cannabis_leaf_morpho_updated). The *x* and *y* coordinates of blade outlines and landmarks were extracted using ImageJ (Abràmoff *et al*., 2004). The outline was extracted using the *wand* tool (setting tolerance to 20 and including “smooth if thresholded” option) and the landmarks were placed using the *multi-point* tool.

Initially, landmarks were placed at the beginning and end of each leaflet, starting from the lower left side, and continuing to the lower right side of the leaf outline. Subsequently, landmarks were placed in the same order on the tips of the leaflets. The final landmark was positioned at the center of the petiolar junction (Fig. **2**, second column). These landmarks delimit the boundaries of the leaflets so that equidistant pseudo-landmarks can later be placed along the contour. The number of landmarks per leaf ranged from 10 to 28, depending on the leaflet number. The raw data containing the coordinates for leaf outlines and landmarks can be accessed on GitHub (https://github.com/BalantM/Cannabis_leaf_morpho_updated).

**Fig. 2.**
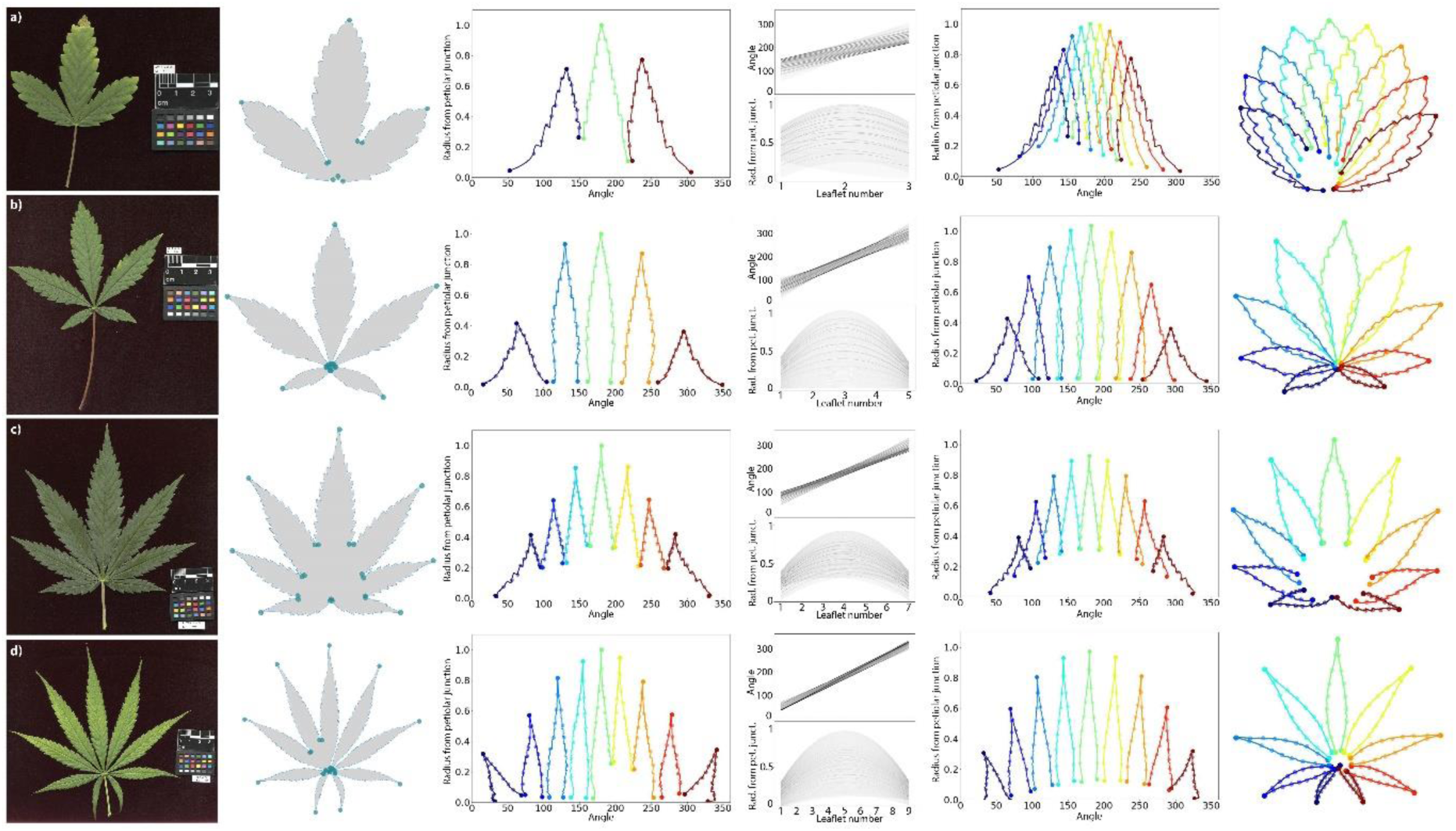
The process of modeling theoretical leaves for a leaf with (a) three leaflets from accession AM15, (b) five leaflets from accession IKL, (c) seven leaflets from accession FUT75, and (d) nine leaflets from accession IK. The first column shows the scans of the leaves, which we use to extract the outline and place the landmarks on the tip, start, and end of each leaflet and on the petiolar junction (second column). These coordinates are used to generate 200 equidistant pseudo-landmarks on each side of each leaflet, sharing the landmark on the tip of the leaflet for a total of 399 pseudo-landmarks. These coordinates are then converted into polar coordinates. Each transformed leaflet is defined with 399 equidistant pseudo-landmarks, with three landmarks, two at the base and one at the tip. Large points are placed every 25 pseudo-landmarks to emphasize that leaflet outlines are defined by points (third column). Second degree polynomials for angles and for radius from petiolar junction are then fitted through these 399 pseudo-landmarks (fourth column). A modeled theoretical leaf with nine leaflets defined by 3,591 pseudo-landmarks can then be modeled using the collection of 798 polynomial models for each leaf (399 polynomial models for angles and 399 for radius from petiolar junction) (fifth column) and visualized in the cartesian coordinate system (sixth column).

### Reconstruction of the new modeled leaves

To analyze leaves with different numbers of leaflets, pseudo-landmarks of each leaflet were modeled as 2^nd^ degree polynomial models of angles and radius as functions of leaflet number within a leaf, in order to use the models to construct a modeled theoretical leaf with a desired number of leaflets. The Python code, presented as a Jupyter notebook with detailed description, is available on GitHub (https://github.com/BalantM/Cannabis_leaf_morpho_updated). The *x* and *y* coordinates of the leaf outline were first interpolated to create an arbitrarily high number of coordinates to increase resolution of the leaf outline. The coordinates of manually selected landmarks were then compared against the high-resolution coordinates of the leaf outline and the nearest neighboring point of the high-resolution coordinates to each original landmark was identified and specified as the new landmark point. Next, the outline and new landmark coordinates were rotated, translated, and scaled so that the central leaflet had a length of one and pointed in the same direction. The transformed points were then interpolated to generate 200 pseudo-landmarks on each side of each leaflet (from the landmark at the bottom until the tip of the leaflet), sharing the landmark on the tip of the leaflet (i.e., a total of 399 pseudo-landmarks per leaflet). These pseudo-landmarks were then converted to polar coordinates, where each point was defined by a radius and angle relative to the landmark of the petiolar junction and tip of the central leaflet (Fig. **2**, third column).

Using the polar coordinates of each leaflet, 2^nd^ degree polynomial models for *x* (angle) and *y* (radius from petiolar junction) values were fit through each of the 399 corresponding pseudo-landmarks for each leaflet using the Python *scipy.optimize.curve_fit* function (Virtanen *et al*., 2020), modeling angle and radius as a function of leaflet number (Fig. **2**, fourth column). Using the coefficients for 2^nd^ degree polynomial models, we then model each pseudo-landmark as a function of leaflet number to reconstruct the new theoretical leaf with an arbitrary number of leaflets. Meaning that for each leaflet, each of the 399 *x* and *y* pseudo-landmarks (i.e., angle and radius from petiolar junction coordinates) was calculated using the 2^nd^ degree polynomial function, with coefficients obtained from the previous step, and the newly defined leaflet number (9 in this case). The optimal number of reconstructed leaflets was tested for the best prediction accuracy in Linear discriminant analysis modeling and the highest accuracy was achieved by reconstructing 9 leaflets (Table **S1**). It is important to note that the reconstructions start with the first real leaflet and end with the last real leaflet. These 9 reconstructed leaflets are then equally divided between these two points.

Nine leaflets were reconstructed using the collection of coefficients of 789 2^nd^ degree polynomial models for each leaf; the 399 models for angle were used to model theoretical *x* (i.e., angle) and 399 models for radius were used to model theoretical *y* (i.e., radius from petiolar junction) pseudo-landmarks as a function of nine leaflets.

The coordinates defining the 3,591 pseudo-landmarks for each of the modeled leaves (399 pseudo-landmarks for each of the 9 reconstructed leaflets) were then plotted and visually inspected. We detected 17 inaccurately modeled leaves, most likely caused by the position of the petiole landmark compared to the landmark marking the start and end landmarks of the leaflet. A total of 341 *Cannabis* leaves were then used in the analysis.

### Validation of the leaf modeling approach

To validate our modeling approach, we extracted the polar coordinates of the original central leaflets (Fig. **3a**) and central leaflets of the modeled leaves (Fig. **3b**) and used them in Procrustes analysis using *Procrustes* function from scipy.spatial module (Virtanen *et al*., 2020). Procrustes analysis minimizes the distance between all points for a set of landmarks/pseudo-landmarks between two samples through translation, rotation, and scaling, and returns new points of the two sets, superimposed to each other (Fig. **3c**). We then calculated the Procrustes distance between the original central leaflet (angle and radius coordinates) to its corresponding modeled reconstruction, a measure of their similarity. The mean distance was calculated and compared to that of simulated bootstrapped mean values by resampling (10,000 resamples) through randomly sorting original leaflet coordinates against coordinates of reconstructed leaflets.

**Fig. 3.**
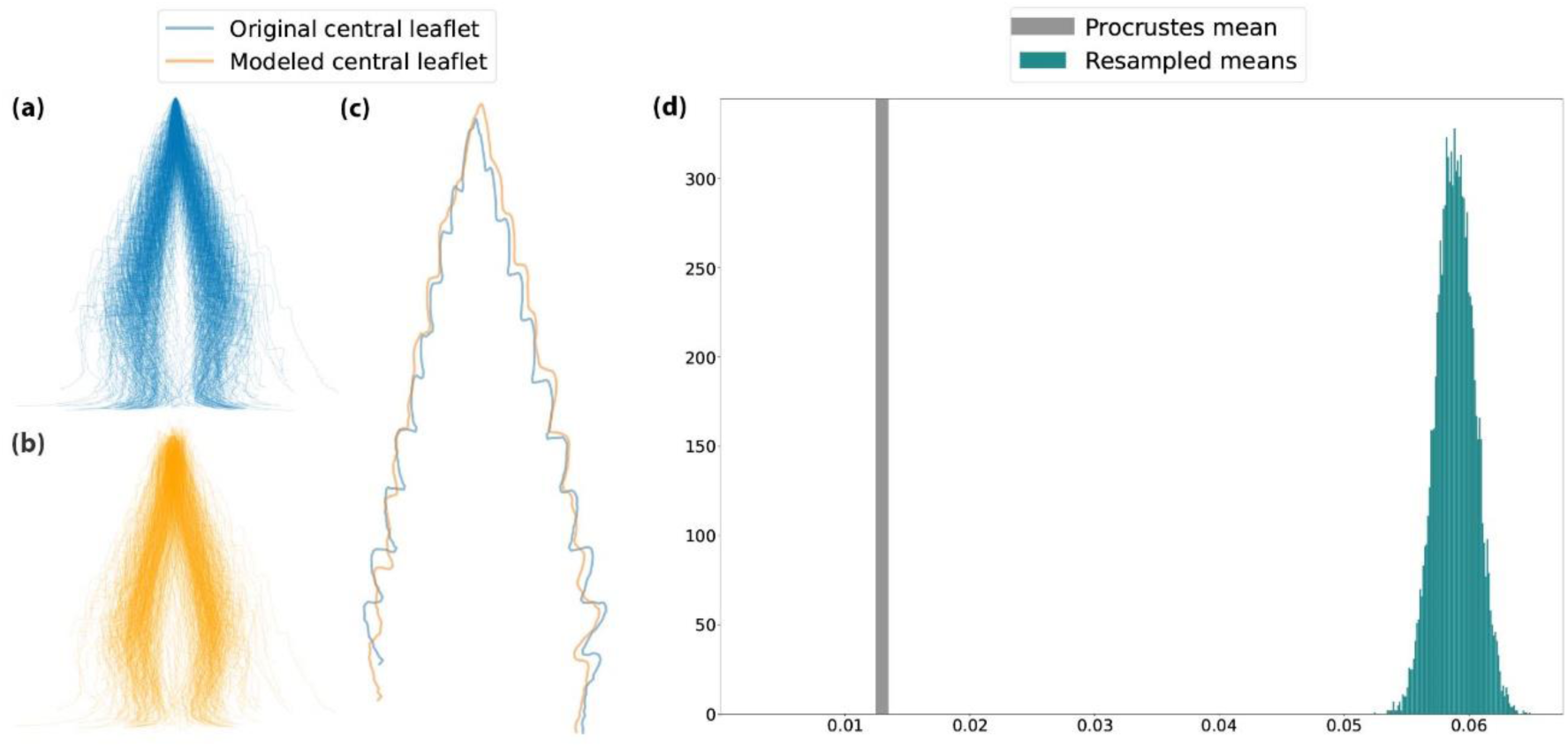
Modeling approach validation using Procrustes analysis and bootstrap resampling. The (a) original and (b) modeled central leaflets in polar coordinate system were superimposed (c) and Procrustes distances calculated. (d) The resampled mean was plotted as a distribution (green histogram) against the actual Procrustes mean (grey vertical line).

### Morphometric analysis of the central leaflet shape using previously established methodologies

The width-to-length ratio (W/L ratio), first described by Anderson (1980), was frequently used to describe the shape of *Cannabis* leaves or even differentiate between different *Cannabis* taxa. With previously established morphometric methods, the shape analysis of central leaflets (that all leaves share) would also be possible, using EFDs or pseudo-landmarks approach. To evaluate the effectiveness of these two previous methods for the shape analysis of *Cannabis* leaves, we first extracted the Cartesian coordinates of central leaflets (Fig. **4a**), that were previously scaled, rotated and translated, so that they were all pointing in the same direction and had the length of one. We then interpolated 200 pseudo-landmarks on each side of each leaflet, sharing the landmark on the tip of the leaflet (i.e., a total of 399 pseudo-landmarks per leaflet).

**Fig. 4.**
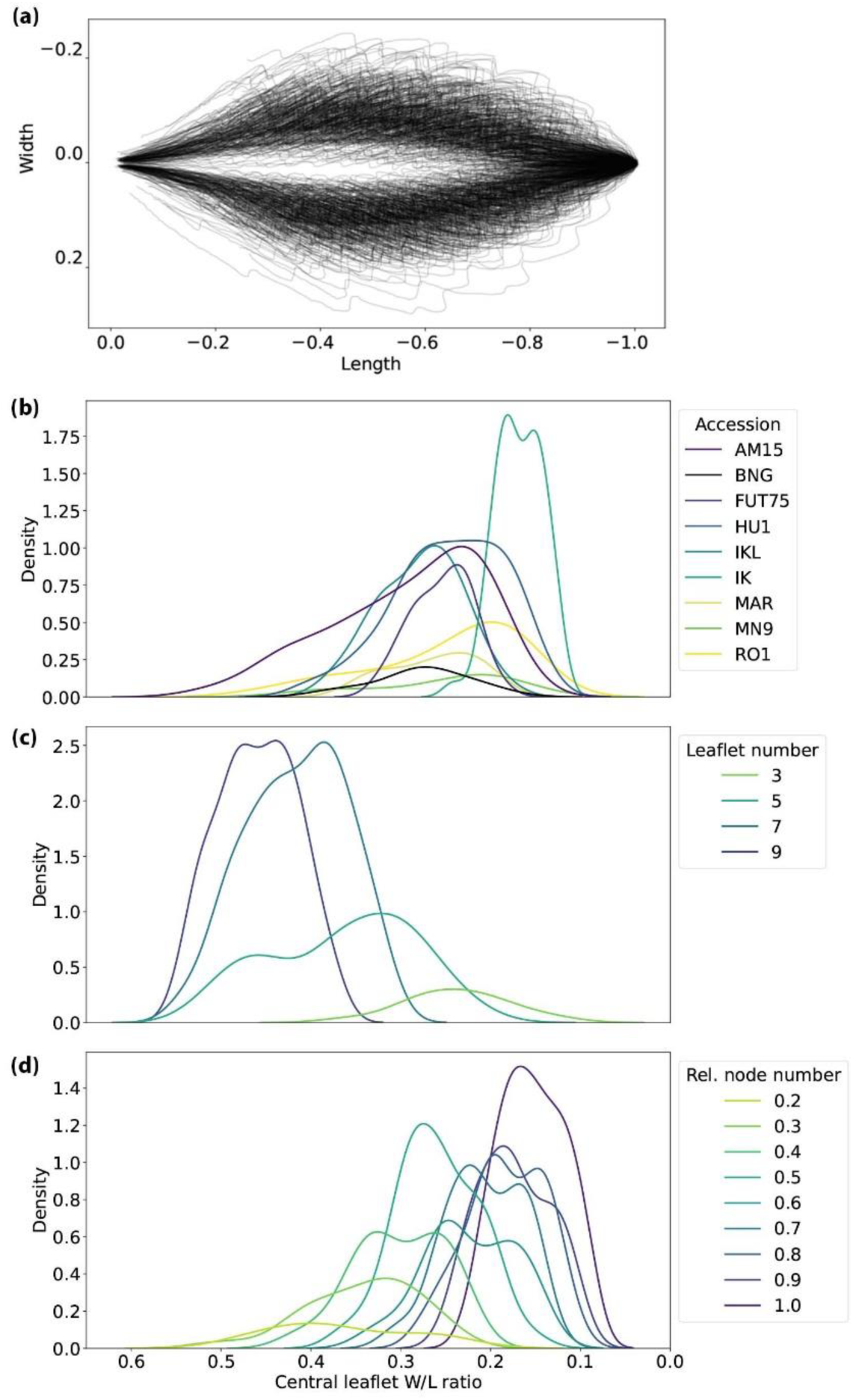
Analysis of leaf shape using the approach adapted from Anderson (1980). (a) Visualization of the 341 central leaflets used in the analysis. W/L ratios plotted by (b) accession, (c) leaflet number and (d) relative node number.

To measure the W/L ratio, we calculated width of the leaf (as the leaves were already normalized to length of one), calculating the minimum bounding rectangle. The distribution of widths was then plotted using Python package *seaborn.kdeplot*. To see if the analyzed accessions differed significantly in their W/L ratios, Kruskal-Wallis test was calculated using *stats.kruskal* function from the scipy.stats module. To see which of the accessions differ in W/L ratio, we calculated Dunn’s Multiple Comparison Test with *scikit_posthocs* package in Python (Terpilowski, 2019), using the *posthoc_dunn* function.

Linear discriminant analysis (LDA) was applied to model accession, leaflet number, and relative node number as the function of central leaflet coordinate values, using the *LinearDiscriminantAnalysis* function from the scikit-learn module in Python (Pedregosa *et al*., 2011). To test the performance of the LDA model, the dataset was divided into two parts. Since most of the analyzed leaves exhibit opposite phyllotaxy, wherein the nodes were represented by two leaves (a and b) in the same developmental phase with the same number of leaflets, the dataset was split into a training dataset (leaf a) comprising 180 leaves and a test dataset (leaf b) containing 161 leaves. The *predict* function from *LinearDiscriminantAnalysis* in the scikit-learn module was used to predict the accession identity, leaflet number, and relative node number, based on the central leaflet coordinate values. The accuracy of the LDA model was calculated and visualized using the function *confusion_matrix* from scikit-learn. Spearman Rank Correlation was calculated for true and predicted results for relative node number with *spearmanr* function from the scipy.stats module.

### Data analysis of modeled leaves

A principal component analysis (PCA) was performed on the coordinates of the modeled leaves using scikit-learn module in Python and proportions of explained variance for each principal component and the cumulative variance was calculated. Points representing the leaves were colored by the accession identity, leaflet number, or relative node number (Fig. **5**). To see which of the first two PCs explains most of the leaf shape variation for accessions, leaflet number and relative node number, Kruskal-Wallis test was calculated using *stats.kruskal* function from the scipy.stats module. To visualize an average leaf for each accession, leaflet number, and relative node number, the average coordinate values of modeled leaves were calculated for each of the categories and plotted using the Matplotlib module in Python (Fig. **5**).

**Fig. 5.**
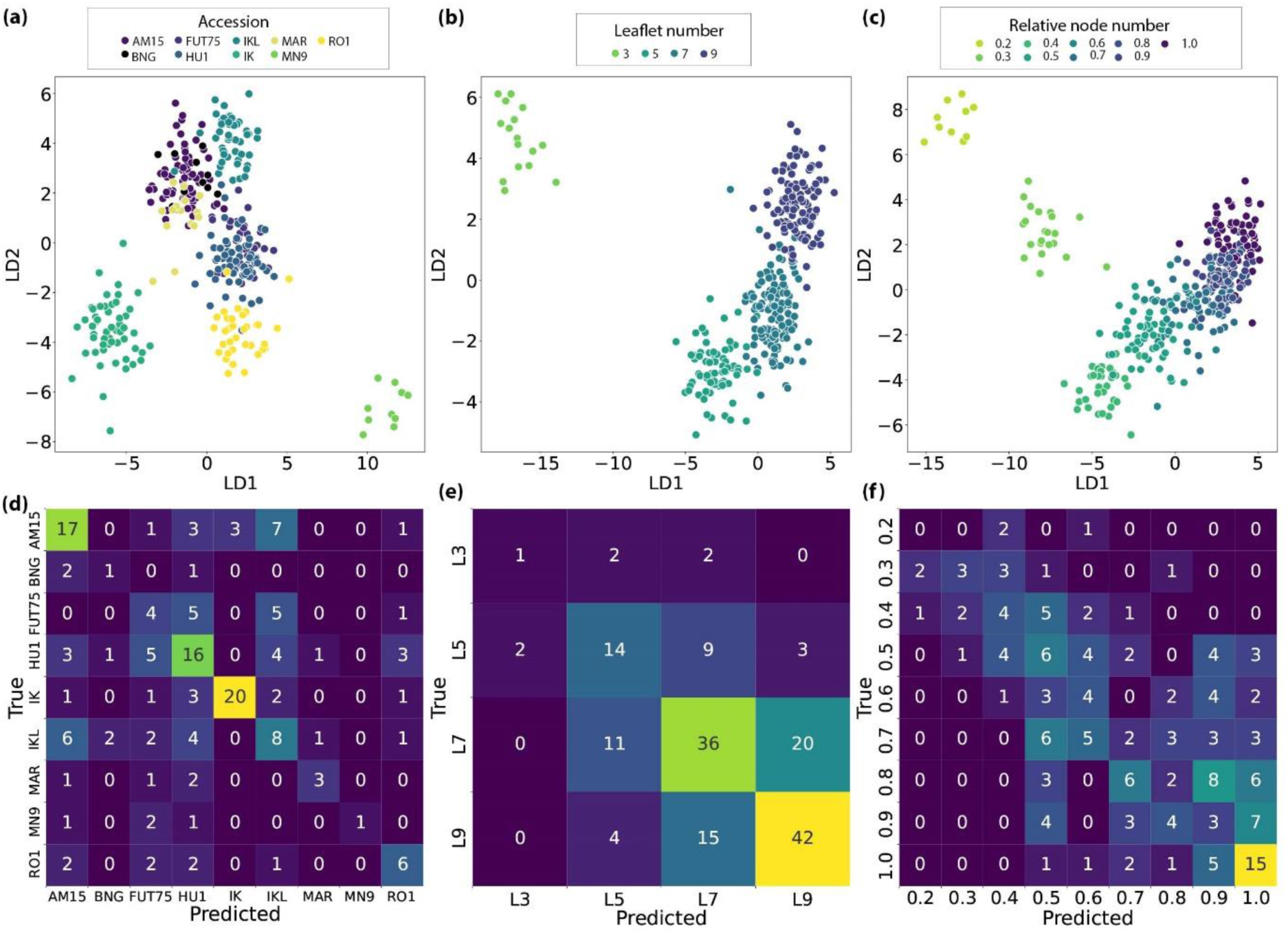
Accession, leaflet number and relative node numbers prediction of *Cannabis* leaves using the outline of central leaflets. Linear discriminant analysis (LDA) plots for (a) accession, (b) leaflet number and (c) relative node number. In the lower row, the confusion matrices show the true and predicted identities for (d) accessions, (e) leaflet number, and (f) relative node number using the LDA model on the split test and train dataset.

To see if the modeled leaves can be used to model accession, leaflet number, and relative node number, we followed the same steps as before for shape analysis of central leaflet. Linear discriminant analysis (LDA) was applied to model accession, leaflet number, and relative node number. The dataset was again split into a training and test dataset to see if we were able to predict accession, leaflet number, and relative node number identity, based on the coordinates of modeled leaves. The same was done on a combined dataset with 3990 coordinates, created by concatenating coordinates of modeled leaves and the coordinates of the original central leaflets.

## RESULTS

### Heteroblastic changes in leaflet number along the main axis

Over 460 *Cannabis sativa* leaves were collected, scanned, and their leaflet number recorded. The leaves exhibited a profound heteroblastic juvenile-to-adult progression along the axis, but the changes were not uniform between the accessions (Fig. **1**). In the few rare cases where the leaves in the lower nodes were present, the first nodes always started with a simple serrated leaf. The second leaf usually had three leaflets and the most frequent leaflet number in the third node was five. However, the leaflet number in the nodes above varied dramatically between accessions. The number of nodes before the transition into the inflorescence in each of the plants also varied. We therefore calculated relative node number, a fractional number between 0 at the shoot base to 1 at the inflorescence transition, to compare the node leaves between plants.

### Validation of the leaf modeling approach

The modeling approach was validated by calculating the mean Procrustes distance of modeled central leaflet coordinates to original central leaflet coordinates using 10,000 bootstrap replicas, assessing resampled means against the actual Procrustes mean value. None of the 10,000 resamples yielded a mean lower than the observed Procrustes value, confirming the robustness of the novel modeling approach (Fig. **3d**).

### Width-to-length (W/L) ratio and central leaflet shape analysis

Our results indicate that the width-to-length (W/L) ratio of central leaflets is not able to differentiate well between different *Cannabis* leaf accessions based on this information alone (Fig. **4**). While the Kruskal-Wallis test did show overall significance between accessions (Table **S2**), Dunn’s post hoc indicated significance in leaf morphology for just one accession (Table **S3**). The W/L ratio significantly differs from the rest only for the IK accession, characterized by particularly narrow leaves (Table **S3**). The Kruskal-Wallis test was also significant for leaflet numbers and relative node numbers (Table **S2**). Dunn’s post hoc revealed that while we can differentiate between leaflet numbers based on the W/L ratio of central leaflet, we can only separate the lower and higher relative nodes (Table **S3**).

To test whether the outline of the central leaflet can better predict the genetic and developmental identity of *Cannabis* leaves, we used Linear discriminant analysis (LDA) to model each factor as a function of 399 pseudo-landmark points defining the shape of central leaflet (Fig. **5a-c**). To evaluate model accuracy, accession was treated as a categorical variable, as was leaflet number, as it not only has a small number of levels (3, 5, 7, and 9 leaflets), but each level is well separated from the others. To evaluate the accuracy of relative node number, we treated it as a continuous variable, due to a high number of levels (9) that continuously overlap with each other. Models revealed low accuracy, as the accession was correctly determined only in 47.20% (Table **2**). The LDA model for the shape of central leaflet showed no overlap for the accessions IK and MN9, but the remaining accessions showed significant overlap (Fig. **5a**). The confusion matrix revealed that only two accessions were correctly identified more than half the time (AM15 – 53.13% and IK – 71.43% prediction accuracy) (Fig. **5d**). The LDA model showed better success when identifying the leaflet number (57.76% overall accuracy) and relative node number, where the true and predicted values show significant, but moderate correlation (rho = 0.629, p < 0.0001) (Fig**. 5b, c, e, f**; Table **2**).

**Table 2.** Predictive power of genetic and developmental identities using the LDA model on the central leaflet shape.

| | Correct<br>prediction [n] | False<br>prediction [n] | Prediction<br>accuracy [%] | Correlation<br>coefficient [$\rho$] | p value |
| --- | --- | --- | --- | --- | --- |
| Accession | 76 | 85 | 47.20 | NA | NA |
| Leaflet number | 93 | 68 | 57.76 | NA | NA |
| Relative node number | NA | NA | NA | 0.629 | < 0.0001 |

### Principal component analysis on modeled leaves (PCA)

Using the outline and landmark coordinates of 341 leaves, we modeled new theoretical leaves, all with nine leaflets. Each leaf is defined by 3,591 pseudo-landmarks, which overcomes the problems associated with variable leaflet numbers and permits dimension reduction using PCA (Fig. **6a-c**) and the visualization of average *Cannabis* leaves (Fig. **6d-f**). The first and second PCs account for 85.85% and 7.25% of the shape variation, respectively (Fig. **6a-c**). Examining the PC1 and PC2 with Kruskal-Wallis test reveals that that accession, leaflet number and relative node number all vary significantly along the first PC axis. The variation along the PC2 for accession and leaflet number is less pronounced, however still significant, while PC2 values for relative node numbers do not vary significantly (Fig. **6**; Table **3**). This indicates that the changes in leaf shape between accessions are not independent from developmental variation. That a facet of variation in accession leaf shape covaries with developmental variation across the shoot in leaflet and relative node number suggests a heterochronic mechanism by which accession differences in leaf shape arise from changes in developmental timing, and contrasts with the historical focus on changes in timing arising from plasticity (Goebel, 1908; Ashby, 1948).

**Fig. 6.**
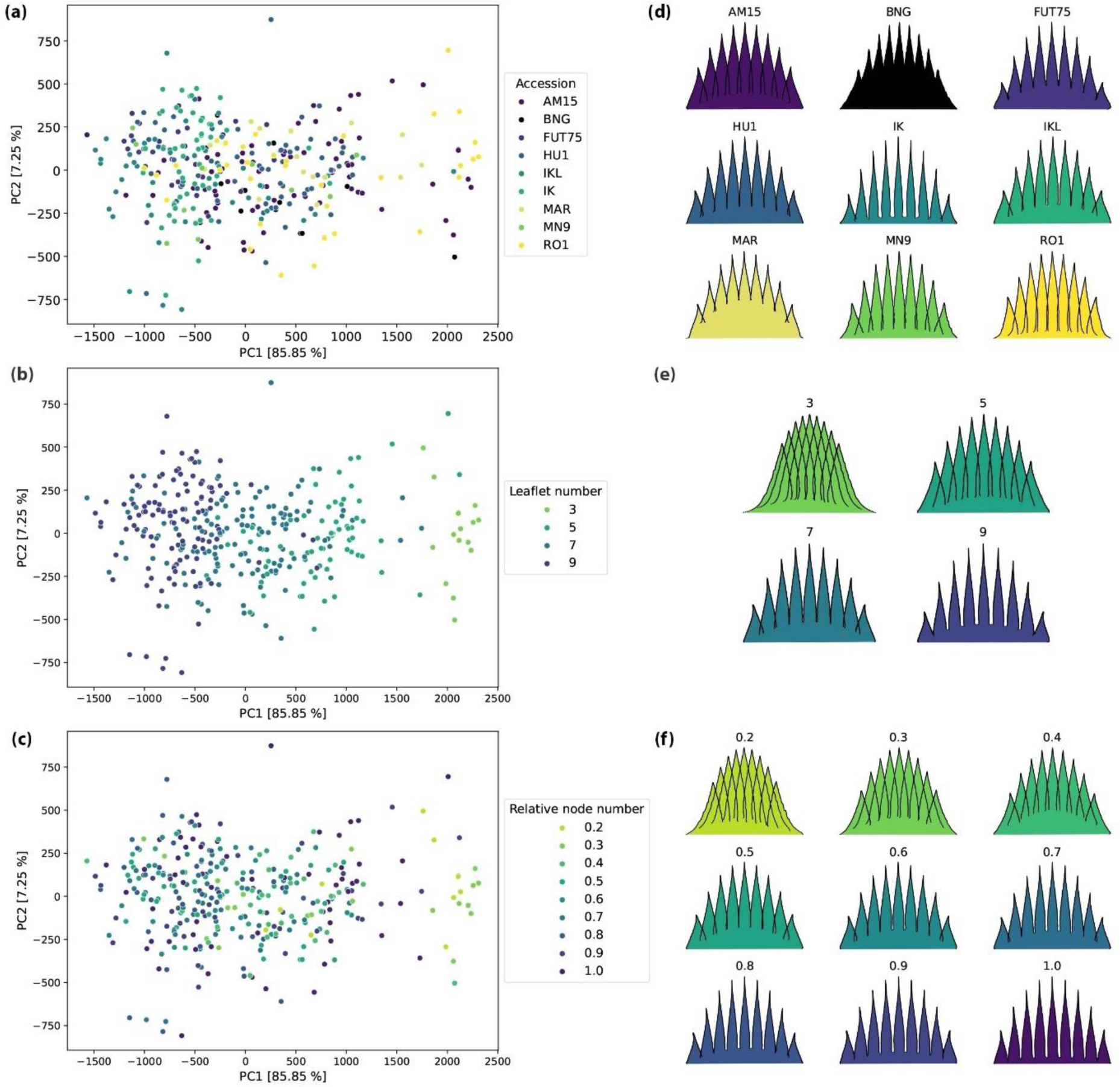
Principal component analysis (PCA) of the accessions performed on modeled leaves using the 3,591 pseudo-landmarks (a-c). The first PC explains 85.58% and the second 7.25% of variation. The images on the right show the average modeled leaf shapes for each of the (d) nine analyzed accessions, (e) leaflet number and (f) relative node number.

**Table 3.**
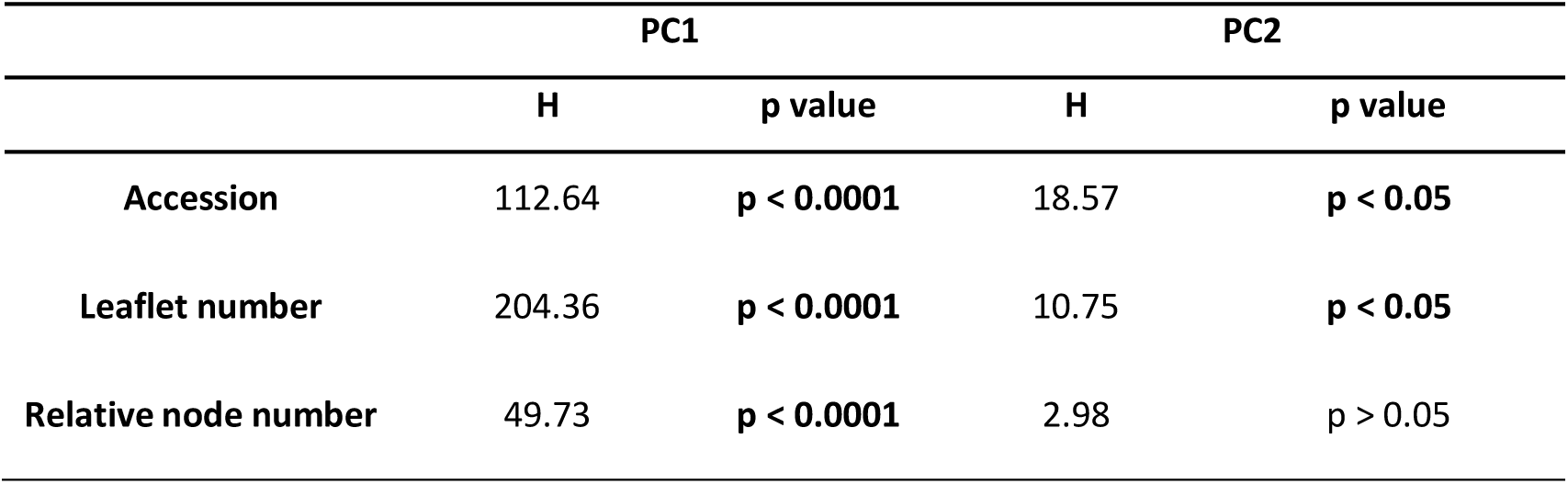
Kruskal-Wallis test was used to test the leaf shape variation along PC1 and PC2 for accessions, leaflet number and relative node number.

|  | PC1 |  | PC2 |  |
| --- | --- | --- | --- | --- |
|  | H | p value | H | p value |
| Accession | 112.64 | $p < 0.0001$ | 18.57 | $p < 0.05$ |
| Leaflet number | 204.36 | $p < 0.0001$ | 10.75 | $p < 0.05$ |
| Relative node number | 49.73 | $p < 0.0001$ | 2.98 | $p > 0.05$ |

The average modeled leaf shapes show that the most pronounced change in leaf shape between the accessions and during the development corresponds to narrow vs. wide leaflets that are stereotypical descriptions of *sativa* vs. *indica* or *wide*- vs. *narrow- leaflet* drug varieties. Furthermore, the leaves with the lower number of leaflets have more acute leaflet tips, that slowly transition into acuminate. Additionally, the outer leaflets in the leaves from lower nodes (and in certain accessions) are longer, compared to the central leaflet, and become shorter higher up (Fig. **6d-e**).

### LDA and prediction of genetic and developmental identities on modeled leaves

As in the analysis of central leaflet shape before, we used LDA to model accession, leaflet number and relative node number as a function of all 3,591 pseudo-landmark points defining the complete modeled leaves (Fig. **7**). Accuracy of the model was calculated on the split dataset, treating accession and leaflet number as categorical and relative node number as continuous variable. LDA models for both accession and leaflet number were highly accurate (73.29% and 99.38%, respectively) (Table **4**), significantly improving the results obtained by analyzing solely the outline of the central leaflet (Table **2**). The model for relative node number is highly accurate as well, as inferred by a highly significant Spearman’s rank correlation coefficient value between actual and predicted values (rho = 0.747, p < 0.0001) (Table **4**).

**Fig. 7.**
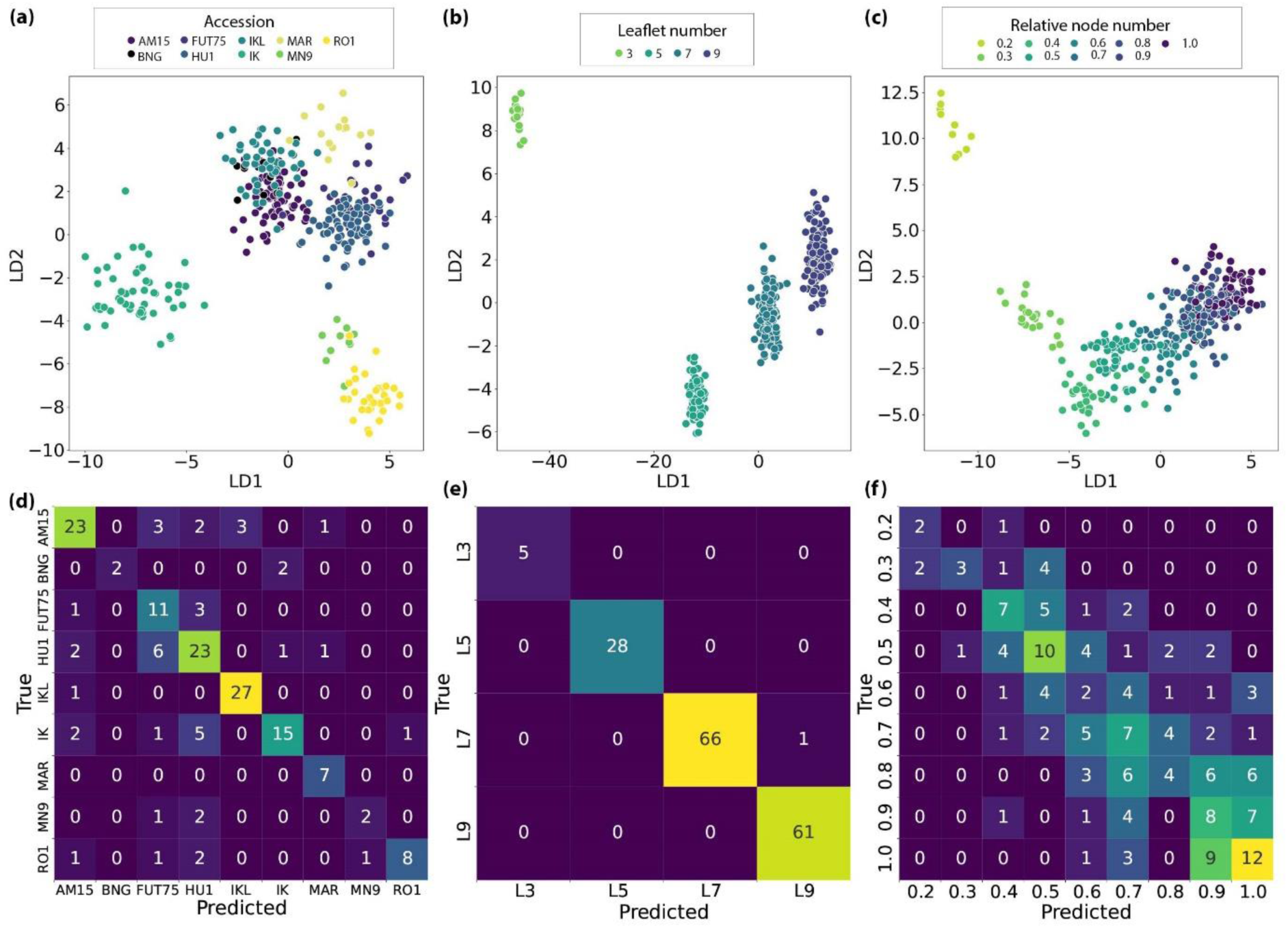
Accession, leaflet number and relative node numbers of *Cannabis* leaves can be predicted independently of each other using modeled leaves. Linear discriminant analysis (LDA) plots for (a) accession, (b) leaflet number and (c) relative node number. In the lower row, the confusion matrices show the true and predicted identities for (d) accessions, (e) leaflet number, and (f) relative node number using the LDA model on the split test and train dataset.

**Table 4.**
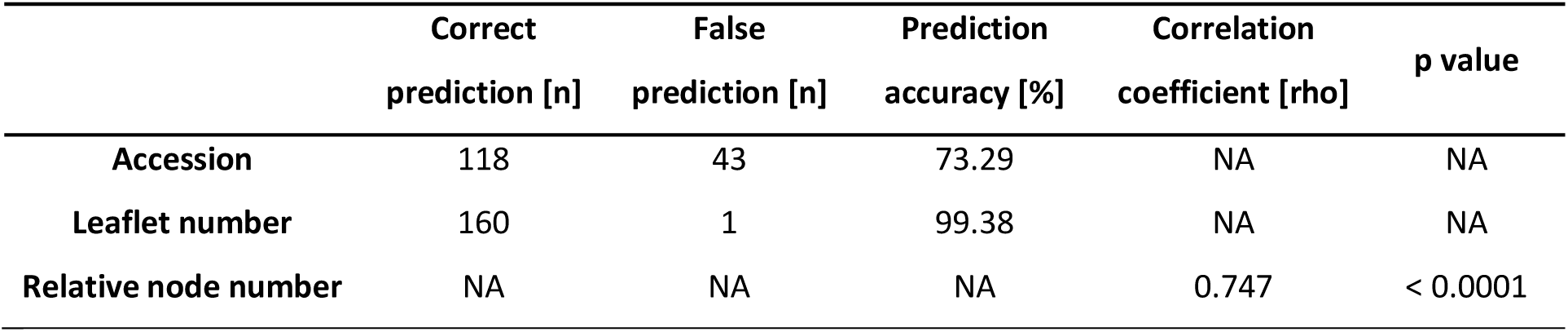
Predictive power of genetic and developmental identities using the LDA model on the modeled leaves.

| | Correct<br>prediction [n] | False<br>prediction [n] | Prediction<br>accuracy [%] | Correlation<br>coefficient [$\rho$] | p value |
| --- | --- | --- | --- | --- | --- |
| Accession | 118 | 43 | 73.29 | NA | NA |
| Leaflet number | 160 | 1 | 99.38 | NA | NA |
| Relative node number | NA | NA | NA | 0.747 | < 0.0001 |

A confusion matrix reveals that the LDA model in most cases had a high accuracy for predicting accession identity (Fig. **7d**; Table **4**), much higher, as compared to the accuracy achieved by using only the outline of the central leaflet (Fig. **5d**, Table **2**). Accessions IK, RO1, and MN9 show practically no overlap in LDA space, while AM15, BNG, FUT75, HU1, IKL and MAR show more overlap (Fig. **7a**). The model showed an almost 100% success rate in determining leaflet number, again, much higher than before.

Results of both methods revealed that leaves with only 3 leaflets are markedly different from the rest, and the prediction model on theoretical leaves consistently classified them correctly (Fig. **7e**). Leaves with 5 to 9 leaflets showed less pronounced differences in shape, resulting in a slightly lower accuracy of the prediction model for these cases. However, an examination of the confusion matrix revealed that misclassifications only occurred once between leaves with neighboring leaflet numbers (7 and 9 leaflets) (Fig. **7e**). The marked difference in shape of leaves with 3 leaflets from the rest may suggest that this developmental mechanism is biased towards variation at the base of the shoot. Similar to leaflet number, the confusion matrix for the relative node model reveals high rates of misclassification between the neighboring relative node numbers, as is expected, and leaves from lower nodes were very rarely classified as those from higher nodes (Fig. **5f**). A pronounced change in leaf shape occurs between the relative nodes 0.3 and 0.4, while the shape changes in later relative nodes are more gradual (Fig. **7c**).

Compared to only using the modeled leaves, the accuracy of the LDA model did not improve significantly when using a combined dataset. A confusion matrix revealed that the LDA model (Fig. **S1**) was slightly less successful in accession identity classification (71.43%) but was higher for leaflet number (100%). The Spearman’s rank correlation coefficient was slightly higher and highly significant (rho = 0.748, p < 0.0001) (Table **5**).

**Table 5.** Predictive power of genetic and developmental identities using the LDA model on a combined dataset.

|  | Correct<br>prediction [n] | False<br>prediction [n] | Prediction<br>accuracy [%] | Correlation<br>coefficient [rho] | p value |
| --- | --- | --- | --- | --- | --- |
| Accession | 115 | 46 | 71.43 | NA | NA |
| Leaflet number | 161 | 0 | 100 | NA | NA |
| Relative node number | NA | NA | NA | 0.787 | < 0.0001 |

## DISCUSSION

Like grapevines, striking variation in leaf shape (Fig. **1**) has historically played a significant role in taxonomic classification of *Cannabis*. Leaf shape and differences in phyllotaxy were among the characters Lamarck used to describe a new *Cannabis* species (Lamarck & Poiret, 1783). Anderson (1980) introduced a quantitative approach by quantifying the length-to-width ratio of the central leaflet. Further studies using different characters—including plant height, stem diameter, achene shape, and phytochemical profiles—to characterize accessions have only confirmed the importance of leaf characteristics (Small *et al*., 1976; Hillig, 2005a). The central leaflet width-to-length ratio has been adopted by researchers as one of the main characters for determining species, subspecies, biotypes and chemotypes of *Cannabis* (Hillig, 2005a; Clarke & Merlin, 2013; McPartland & Small, 2020). However, this method is only able to capture a limited aspect of leaf shape variation, neglecting other important characteristics that we measure in this study, such as leaflet outlines, serrations, angles, and relative changes in leaflet shape across the leaf. By modeling leaflet shape as a function of leaflet number, we model theoretical leaves with the same number of leaflets for which high densities of corresponding pseudo-landmarks capture high resolution shape features (Fig. **2**). To validate the modeling approach, we have compared the outline of the original central leaflet and the outline of the modeled theoretical central leaflet. The Procrustes analysis showed that the two leaflets are very similar in shape, and that the modeling is even able to preserve the serration pattern to some degree (Fig. **3c**). The modeling approach validated using 10,000 bootstrap replicas confirmed the robustness of the novel modeling approach (Fig. **3d**). This method can be applied not only on palmately composed leaves as in *Cannabis* but is also possible to use on pinnate and lobed leaves. To demonstrate the proof of concept, we applied the method to a pinnate leaf of *Cardamine flexuosa* With. and lobate leaf of *Quercus macrocarpa* Michx. (Fig. **8**), showing the method could be applied in other leaf types. However, the method needs to be improved before being applied to other species but shows the possible utility of intra-leaf modeling.

**Fig. 8.**
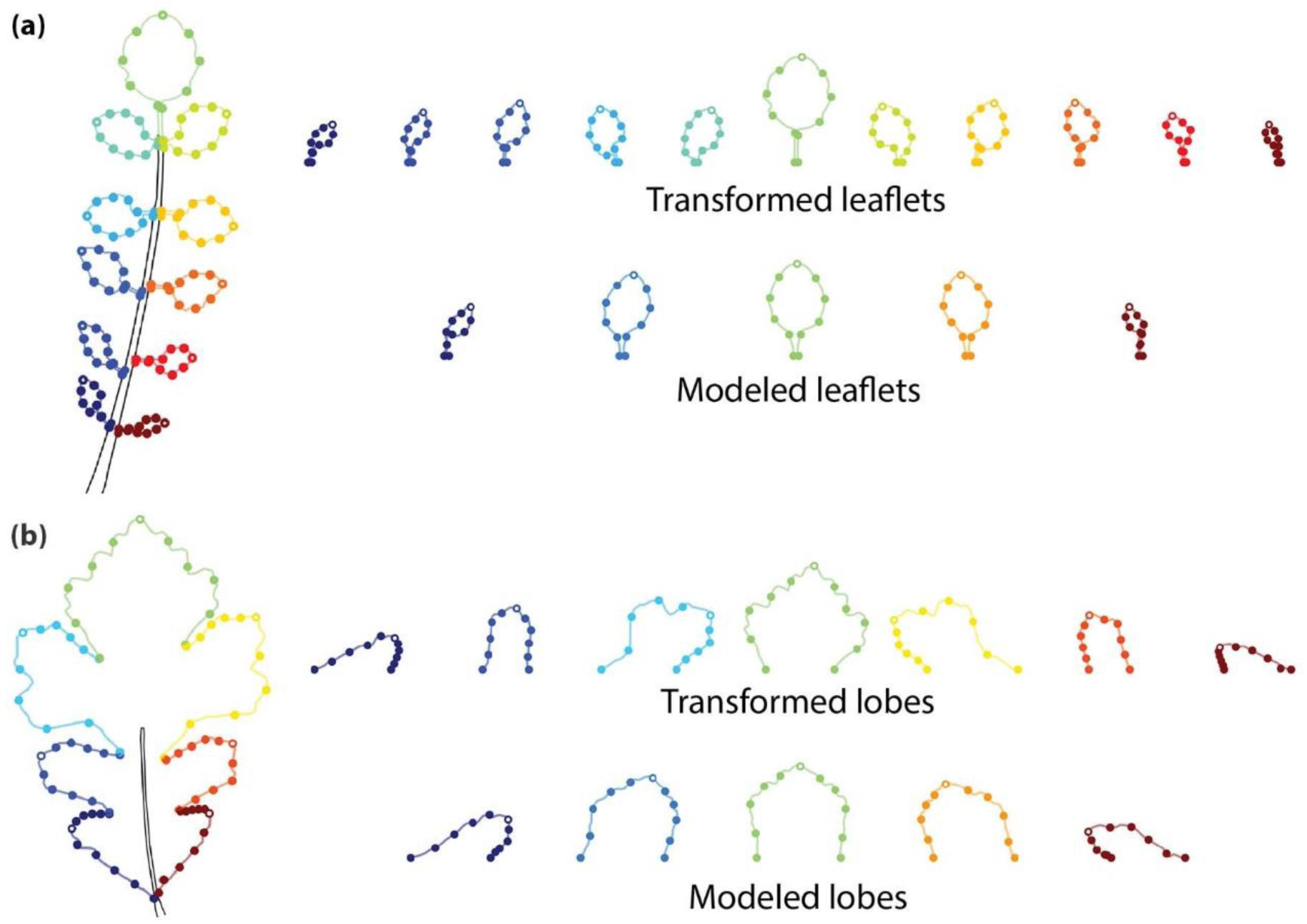
Intra-leaf modeling of leaflets and lobes extended to pinnate leaves: Leaves from (a) *Cardamine flexuosa* and (b) *Quercus marcocarpa*. Leaflets and lobes are defined by 100 equidistant pseudo-landmarks on each side, each defined by three landmarks, two at the base and one at the tip. Large points are placed every 20 pseudo-landmarks to emphasize that leaflet outlines are defined by points. The landmarks defining the base of each leaflet or lobe are aligned to the rachis or midvein and the transformed leaflets and lobes have been oriented parallel to the rachis, as defined by the landmarks at their base. The modeled leaflets and lobes are created from 2^nd^ degree polynomial models for each *x* and *y* coordinate value for each pseudo-landmark as a function of leaflet or lobe number. From these models, an equivalent number of modeled leaflets or lobes can be reconstructed (in this case, five), permitting morphometric analysis.

The method presented in this study can accurately determine accession based on leaf shape, regardless of its developmental stage (Fig. **7a, d**). The method not only works effectively on stabilized or cloned cultivar accessions but also on wild or feral accessions cultivated from seed that can exhibit distinct plant phenotypes (Table **1**), indicating its robustness and potential value in future germplasm classification. Compared to the low accuracy and prediction ability of the previously known methods (W/L ratio and shape analysis of central leaflets), the newly proposed method demonstrates significantly improved results (Table **2**, 4**, S2, S3**). The combined dataset of both, data for modeled leaves and outline of the central leaflet, did not return significantly better results, further confirming the effectiveness of the new modelling approach.

When observing the shape changes between averaged leaves for accessions and between developmental stages, the most obvious are changes in leaflet widths, similar to stereotypical classifications of *sativa* and *indica* plants or *wide*- vs. *narrow- leaflet* drug varieties. However, other important changes in shape occur, such as transition from acute to acuminate leaflet tip and changes in the relative length of outer most leaflets compared to the central leaflet, that previous methods could not successfully capture (Fig. **6d-f**). The reliance on the non-quantitative leaf shape descriptors in previous methods has led to numerous cultivars with unreliable names, inconsistent genetic origins, and phytochemical profiles (Sawler *et al*., 2015; Schwabe & McGlaughlin, 2019; Jin *et al*., 2021a; Watts *et al*., 2021). For example, Jin *et al*. (2021b) conducted a study on clones of 21 cultivars and found a strong negative correlation between the width and length ratios of central leaflets and CBD, and a positive correlation with THC; however, Vergara *et al*. (2021) and Murovec *et al*. (2022) were unable to confirm these findings. All three studies used low-resolution morphometric approaches. Sex of the plants also plays a crucial role in the cannabis industry, where the presence of male plants and inevitable pollination leads to decreases in cannabinoid production as plants shift the use of energy into seed development. Several methods have been employed to differentiate between male and female plants at early stages, but only genetic methods were successful so far (Toth *et al*., 2020; Prentout *et al*., 2020; Campbell *et al*., 2021; Balant *et al*., 2022; Torres *et al*., 2022). Our results quantify the variation in leaf shape between accessions that can potentially be used to classify accessions and predict chemical profiles and plant sex faster and more accurately.

Unlike grapevine, where developmental variance is orthogonal and separate from genetic variance, in *Cannabis* these two factors are correlated. That the developmental source of variation is colinear with accession identity suggests that part of the differences between accession leaf shape is explained by shifts in developmental timing, or heterochrony. *Cannabis* plants demonstrate extreme phenotypic plasticity depending on the environmental conditions in which they grows (Small, 2015). Some *Cannabis* accessions are photoperiod dependent and can remain in vegetative phase for longer periods of time under long-day conditions (typically 18h darkness and 6h light), until the transition to short-day (12h of darkness and 12h of light) induces the formation of the apical inflorescence. Previous investigations showed that other morphological changes, such as decrease in leaf area, number of leaflets per leaf and serration number, occur after the change in the environmental conditions one or two nodes after (Heslop-Harrison & Heslop-Harrison, 1958; Hesami *et al*., 2023). However, differences, especially in flowering time and growth rates between cultivars have been observed before (de Meijer & Keizer, 1996; Hillig, 2005a; Spitzer-Rimon *et al*., 2019; Carlson *et al*., 2021; Naim-Feil *et al*., 2021; Stack *et al*., 2021; Chen *et al*., 2022) and differences in cannabinoid profiles, leaflet index and phenological development were proposed as characteristics to discriminate between them (de Meijer & Keizer, 1996). Heterochronic shifts are apparent in the differential rates in which accessions increase leaflet number across nodes, as well as maximum and average leaflet counts across accessions (Fig. **1**). Remarkably, stages in developmental timing are conserved despite being shifted. For example, a significant shape change exhibited between the leaves with 3 and leaves with 5 leaflets, with leaflets becoming more acuminate and narrower. In contrast, changes in shape between leaves with a higher number of leaflets were more gradual. Additionally, we observed a similar shift in leaf shape between the nodes 0.3 and 0.4, potentially indicating a transition between the juvenile and adult phases of leaf development. Similar results were obtained in previous research. Spitzer-Rimon *et al*. (2022) demonstrated that flowering buds were initiated at node 7, while Moliterni *et al*. (2004) analyzing a different cultivar, found developing flower buds in the 4th node, suggesting that transitions in growth phases are conserved but not synchronized across cultivars. Due to the differences in developmental timing between accessions, the use of continuous models along the shoot could further improve the success predicting accession identity, as was the case in grapevine (Bryson *et al*., 2020).

### Conclusions

In grapevine, leaf shape has long been utilized for variety identification. However, in the case of *Cannabis*, previous attempts were hindered by the variability in leaflet numbers. In this study, we present a pioneering method that successfully addresses this issue. By generating theoretical leaves with customizable leaflet counts, we can now employ high-resolution morphometric techniques to accurately classify different wild/feral and cultivated *Cannabis* accessions. Through the use of 3,591 densely placed pseudo-landmarks, we were able to predict the accession identity with almost 74% accuracy. The method works well not only on stabilized cultivars, but also on phenotypically more variable wild/feral accessions grown from seed. Unifying the number of leaflets allowed us, for the first time, to make comparisons among several leaves along the main axis, enabling us to investigate developmental changes in leaf shape and detect heterochronic mechanisms influencing the leaf shape in *Cannabis*. The implications of this new high-resolution method in both the cannabis industry and research extend beyond its role in determining *Cannabis* accessions. It also offers a promising tool for developmental studies, and for studying the correlation between leaf shape and phytochemical profiles and the sex of the plants, where lower-resolution methods provided inconclusive results so far. The method presented here offers a fast, effective, robust, and low-cost tool that can aid the future classification of *Cannabis* germplasm. Furthermore, the use of this methodology extends beyond *Cannabis*, and can be applied to numerous other plant species with palmate, pinnate, and lobate leaves with varying numbers of lobes and leaflets where the use of geometric morphometrics methods was not previously possible to this extent.

## ACKNOWLEDGEMENTS

This research was supported by projects WECANN (CGL2017-84297-R, Ministerio de Ciencia, Innovación y Universidades), Generalitat de Catalunya (grant number 2021SGR00315) and M. Balant FPI predoctoral contract of the Ministerio de Ciencia, Innovación y Universidades (PRE2018-083226). This work is also supported by NSF Plant Genome Research Program awards IOS-2310355, IOS-2310356, and IOS-2310357. This project was supported by the USDA National Institute of Food and Agriculture, and by Michigan State University AgBioResearch. We would like to thank Joan Uriach Marsal from Uriach Laboratories for additional financial support. We would also like to acknowledge Cannaflos—Gesellschaft für medizinisches Cannabis mbH for providing seed of accessions IK, IKL and MAR analyzed in the study, Carlos Sáez, Carlos Ribelles, Airy Gras, Joan Vallès, Magsar Urgamal, Shagdar Tsooj, Marine Oganesian and Nina Stepanyan-Gandilyan for help with the sample collection and the cultivation process, and Paula Bruna for helpful advice on improving the quality of scanned images.

## COMPETING INTERESTS

None declared.

## AUTHOR CONTRIBUTIONS

MB and DHC conceived the study, with OH, TG and DV inputs in a preliminary design phase of the project. MB, TG and DV cultivated the plants. MB and OH selected the method for imaging *Cannabis* leaves. MB and TG scanned the leaves used in the study. MB and DHC developed the morphometric method and MB analyzed the data. MB and DHC wrote the first draft of the paper, that all authors read, commented, and edited.

## DATA AVAILABILITY

The datasets and code for the method developed here are freely available on GitHub (https://github.com/BalantM/Cannabis_leaf_morpho_updated, https://github.com/DanChitwood/pinnate_leaf_modeling).

## SUPPORTING INFORMATION

The datasets and code generated and analyzed in this study are available on GitHub (https://github.com/BalantM/Cannabis_leaf_morpho_updated, https://github.com/DanChitwood/pinnate_leaf_modeling).

## Notes

https://github.com/BalantM/Cannabis_leaf_morpho_updated

https://github.com/DanChitwood/pinnate_leaf_modeling

### Competing Interest Statement

The authors have declared no competing interest.

### Summary of Updates

The title and author affiliations have been updated. Modelling code in the section Reconstruction of the new modeled leaves has been modified and Validation of the leaf modeling approach and Morphometric analysis of the central leaflet shape using previously established methodologies has been added.

