## Supplementary material for "Intra-leaf modeling of *Cannabis* leaflet shape produces leaf models that predict genetic and developmental identities": Fig. S1

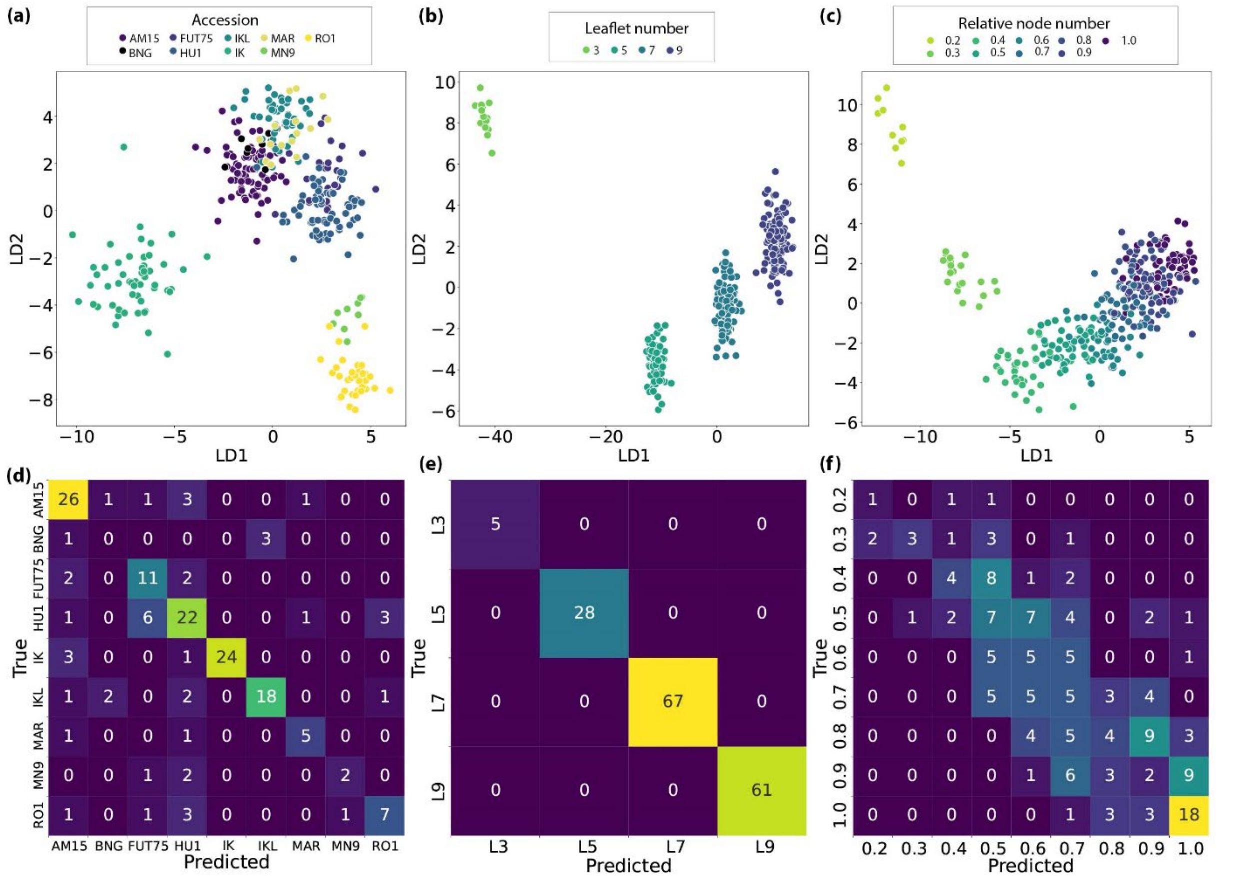


**Fig. S1** Accession, leaflet number and relative node numbers prediction of *Cannabis* leaves using the combined dataset. Linear discriminant analysis (LDA) plots for (a) accession, (b) leaflet number and (c) relative node number. In the lower row, the confusion matrices show the true and predicted identities for (d) accessions, (e) leaflet number, and (f) relative node number using the LDA model on the split test and train dataset.
